# Guide RNA Truncation as a Tunable Approach for Modulating Base Editing Outcomes

**DOI:** 10.64898/2026.09.10.749999

**Authors:** Anila George, Kirti Prasad, Poornasree Sadanandan, Srujan Marepally, Saravanabhavan Thangavel, Kumarasamypet M Mohankumar

## Abstract

Base editing enables programmable nucleotide conversion but can be limited by bystander editing and unintended genomic alterations. Here, we investigated guide RNA (gRNA) truncation as a strategy to modulate base editing outcomes. On editing across multiple genomic loci using ABE8e in HUDEP-2 cells, we found that truncated gRNAs retained measurable editing activity at lengths as short as 15 nucleotides, although activity depended strongly on target sequence, gRNA length, and delivery method. Importantly, truncation altered editing efficiencies at individual nucleotides within the editing window, enabling selective reduction of bystander editing while preserving on-target conversion at several loci, including a therapeutically relevant HbE target. Truncated gRNAs also eliminated detectable large deletions at homologous globin loci, consistent with loss of Cas9 nickase activity at short gRNAs lengths. Similar length dependent effects were observed with cytosine base editing, PAM relaxed ABE8e, and in primary hematopoietic stem and progenitor cells (HSPCs), although the optimal gRNA length varied between contexts. Together, these findings establish gRNA length as an additional parameter for tuning nucleotide level base editing outcomes and reducing bystander editing.

## Introduction

Base editors, particularly the adenine base editor (ABE8e), have emerged as promising tools for precise genome modification with minimal unintended consequences, making it particularly attractive for therapeutic applications(Komor *et al*., 2016; Gaudelli *et al*., 2017). The ability to convert single nucleotides without generating InDels, makes it an attractive tool not only for correction of gene mutations but also for precise gene regulation(Porto *et al*., 2020; Mawson *et al*., 2026). However, certain challenges including bystander editing and the occasional formation of unintended InDels or large deletions, especially in regions with sequence homology, limit the application of base editing. These undesired events can alter the phenotypic outcomes in a bulk-edited population, thereby compromising the utility of base editors(Ravi *et al*., 2022; George *et al*., 2024; Lee *et al*., 2025).

Modulating the precision of base editing is an actively researched area and several groups including us have explored strategies, from enzyme engineering to delivery methods and gRNA modifications to achieve this (Gao *et al*., 2025). By enzyme engineering, researchers have developed modified base editors with narrower editing windows, relaxed PAM requirements and enhanced processivity for base conversion(Tan *et al*., 2019; Richter *et al*., 2020; Tu *et al*., 2022; Li *et al*., 2025; Perrotta *et al*., 2025). However, even with PAM relaxed base editors having narrow editing window, it is difficult to achieve precise conversion, and targeting multiple sites would often require different base editors.

Truncated guide RNAs (Tru-gRNAs) have been extensively studied in the context of CRISPR– Cas9 nucleases, both to understand the mechanisms of Cas9 cleavage as well as to improve specificity. In addition to loss of nuclease activity when truncated below 15 bases, several studies have shown that modest gRNA shortening improves specificity while preserving on-target activity(Fu *et al*., 2014; Zhang *et al*., 2016; Li *et al*., 2023; Jeong, Lee and Lee, 2024). This has been attributed to the differences in the DNA unwinding at the target site as well as conformational changes in Cas9(Josephs *et al*., 2015; Cofsky *et al*., 2022; Sinan and Russell, 2022).

Taking cues from the reports that gRNAs when truncated, retain the ability to bind the target site but unwind DNA differently and abolish strand breaks, we sought to investigate whether modulating the length of gRNAs could mitigate some of the challenges of base editing while preserving editing efficiency. We hypothesised that truncating the gRNA would limit the bases accessible to base editor in the unwound DNA thus modifying the bystander edit. Using HUDEP-2 cells (an immortalized human umbilical cord blood-derived erythroid progenitor cells) and HSPCs (hematopoietic stem and progenitor cells), we comprehensively evaluated the effects of gRNA truncation on base editing efficiency, bystander editing, and formation of large deletions across several therapeutically relevant loci including the *HBG*, *HBB, CCR5* and *CD33*. Our data suggest that tru-gRNAs as short as 15 nucleotides can retain base editing activity in a target-dependent manner. Importantly, truncation of gRNAs resulted in varying editing efficiency at different base positions, even with PAM relaxed base editors thus expanding the scope for modulating bystander effects. Our results also support the potential application of tru-gRNAs in SNP-sensitive genomic contexts and reinforce their potential for complementary use with approaches such as dead Cas9-base editors for safer therapeutic editing.

## Materials and methods

### Plasmid constructs and cloning of the gRNA

gRNA sequences were either selected from previous publications or manually designed. Truncated versions of gRNAs were prepared by sequentially removing the 5’ nucleotide(from the PAM distal end) (Supplementary Table 1). Lentiviral gRNA constructs were prepared as per the previous protocol(Ravi, George and Mohankumar, 2023) and cloned into pLKO5.sgRNA.EFS.GFP/RFP (Addgene #57822) encoding GFP reporter and ampicillin-resistant gene.

For preparation of stable cell lines of base editors (ABE7.10 (Addgene-plasmid#112675), plenti_TADCBEd_puro (constructed from Addgene-plasmid#193835), pLenti-ngABE8e-puro (constructed from Addgene-plasmid#138491)and pLenti-ABE8e-puro (constructed from Addgene-plasmid#138495) plasmids were used.

### Cell culture

HEK 293T cells were cultured in Dulbecco’s modified eagle medium (DMEM) with 10% Fetal Bovine Serum and 1X Pen-Strep. HUDEP-2 cells were cultured in StemSpan SFEM II supplemented with 50 ng/ml SCF (ImmunoTools), 3 U/ml EPO(Zyrop 4000 IU injection), 1× Pen-Strep (Gibco), 1 μM dexamethasone (Alfa Aesar), 1 μg/ml doxycycline (Sigma-Aldrich), and 1× L-glutamine 200 mM (Gibco).

Healthy donor PBMNCs (G-CSF mobilized) were obtained with informed consent under Christian Medical College, Vellore IRB approval. CD34+ cells were isolated by Lymphoprep density centrifugation, RBC lysis, and EasySep positive selection (STEMCELL Technologies), then expanded in HSC media as described (Genovese *et al*., 2014).

### Lentivirus production and transduction

Lentivirus for stable cell lines and for gRNAs were prepared in HEK293T cells using Fugene HD as described previously(Ravi, George and Mohankumar, 2023). The second-generation packaging constructs pMD2.G and psPAX2 (Addgene #12259, 11260) were a gift from Didier Trono. The viral supernatant at 48hrs and 72hrs from 10 cm dish was collected, concentrated, and resuspend in 240ul of 1x PBS separately. From this 30ul was used to transduce 0.1 million HUDEP-2 cells taken in 24 well plate with 1% HEPES and 3μg polybrene (Sigma-Aldrich). Spinfection was carried out at 800G for 30 minutes at room temperature. Stable cell lines were prepared by maintaining the transduced cells in Puromycin. gRNA transduction efficiency was evaluated by measuring GFP expression using flow cytometry and all cells had more than 95% transduction efficiency.

### In Vitro transcription of base editor mRNA

ABE8e and CBETAD mRNA were synthesized from Addgene #138495 and Addgene#193835 using HiScribe® T7 mRNA Kit with CleanCap^®^ Reagent AG(NEB) as per the manufacturer’s protocol. Complete substitution of N-methyl pseudouridine (Trilink) was done and co-transcriptional capping was performed using and Cleancap AG. mRNA was made into aliquots and stored in –80 until use.

### Nucleofection

WT HUDEP-2 and CD34+ cells were expanded in respective media for 48 hours prior to electroporation. Cells (0.5-1 × 10⁶) were pelleted, resuspended in 19 µL MaxCyte buffer (Hyclone) containing 5 µg base editor mRNA and 100 pmoles sgRNA (Synthego), and electroporated using the ‘HSC-3’ program in OC25 × 3 MaxCyte cuvettes. Following electroporation, cells were recovered for 20 minutes (37°C, 5% CO₂) before transferring to and expansion in their respective media for 48 hours.

### DNA isolation and analysis of editing outcomes

Genomic DNA was isolated from the cells on day 6 post transduction using NucleoSpin Blood Mini kit (Macherey-Nagel) as per manufacturer’s protocol. Target loci were amplified with appropriate primers (Supplementary Table 2) using GoTaq Hot Start Master Mix (Promega) and sequenced on the MiSeq platform (Illumina). Editing outcomes were analysed using CRISPResso2(Clement *et al*., 2019).

### Validation of *HBG1-HBG2* and *HBB-HBD* large deletion

Deletion specific primers (Supplementary Table 2) were designed to amplify fusion gene products. For *HBG1-HBG2* deletions, forward primers targeting *HBG2* and reverse primers targeting HBG1 were used. For *HBB-HBD* deletions, forward primers specific to the *HBB* region and reverse primers targeting downstream *HBD* sequences were employed. PCR amplification was performed using GoTaq HotStart Master Mix (Promega) as per manufacturer’s protocol. Products were resolved on agarose gels, confirming deletion-specific bands of approximately 1690 bp (*HBG1-HBG2*) and 788 bp (*HBB-HBD*).

### Statistics

All experiments were performed in independent triplicates unless specified otherwise. Data are represented as mean ± SD.

## Results

### ABE8e can mediate base editing with truncated gRNAs as short as 15 nucleotides

To assess the impact of gRNA length on base editing efficiency, we selected a previously validated gRNA targeting BCL11A binding site (HBG_G2) in gamma-globin promoter (Ravi *et al*., 2022). gRNAs of progressively decreasing length with 5’ truncations (till 10 nucleotides) were designed and cloned in lenti-viral vector and transduced in HUDEP-2 cells stably expressing ABE8e to evaluate base editing efficiency. (**Figure 1A and B**)

**Figure 1:**
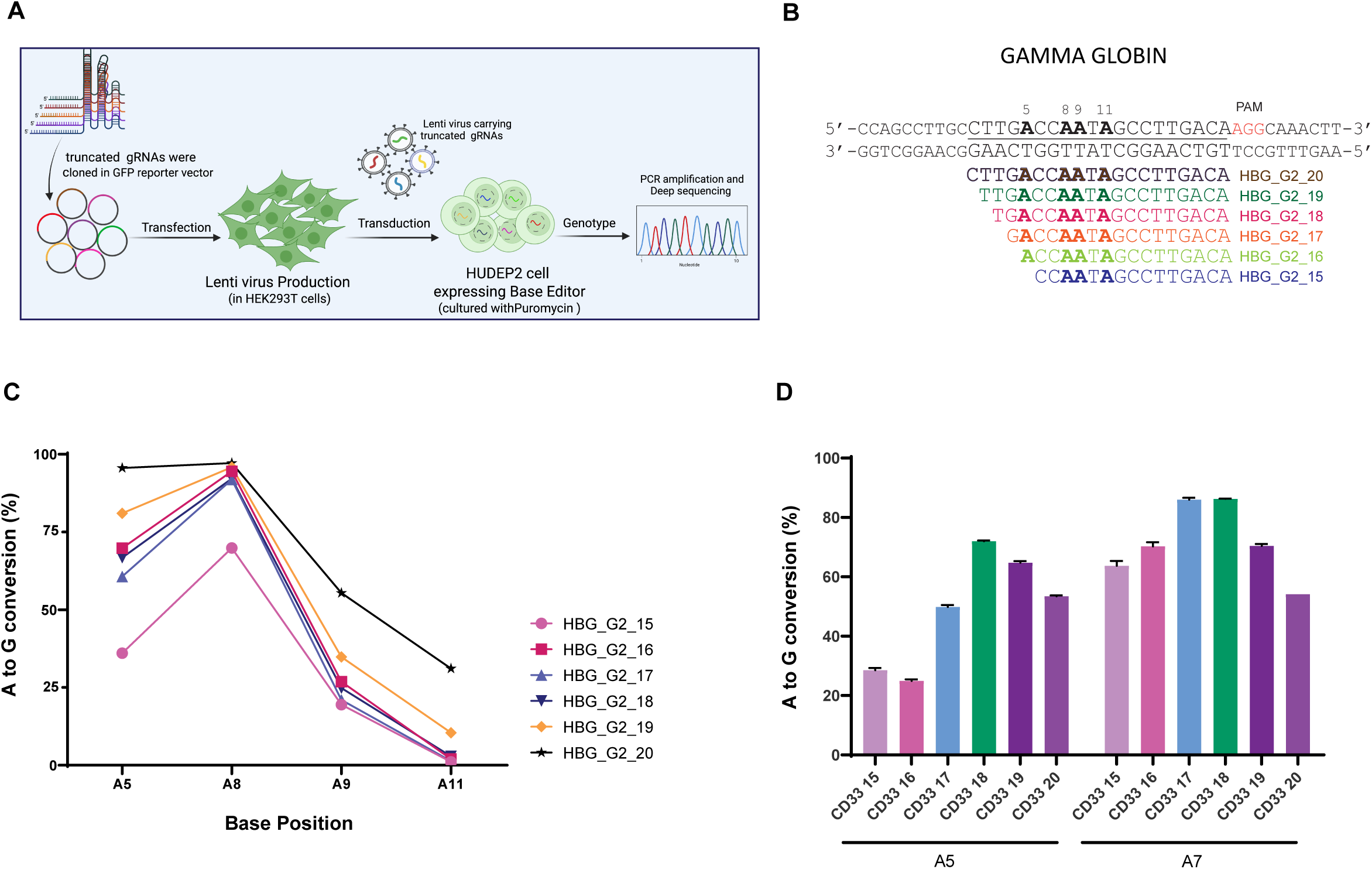
5’ Truncated gRNAs enable efficient base editing in HUDEP-2 cells. A) Schematic representation of the methodology used for testing gRNA efficiency in HUDEP-2 cells B) Sequence of full length and truncated gRNAs targeting the gamma globin locus. Target Adenosines within the editing window are represented in bold letters C) Base conversion efficiency at different positions for 5’truncated gRNAs using ABE8e in the gamma globin locus in HUDEP-2 cells by lentiviral delivery. D) Base conversion efficiency at different positions for 5’truncated gRNAs in the CD33 locus in HUDEP-2 cells using ABE8e by lentiviral delivery. Data from three independent replicates are plotted as mean± SD.

At this locus, progressive 5′ truncation generally reduced editing efficiency, with detectable editing retained till 15 nucleotide gRNA (**Figure 1C**) but further truncation resulted in a complete absence of editing (data not shown). As it has previously been shown that Cas9 loses its nuclease activity with 15 nucleotide gRNA or lower(Fu *et al*., 2014), we sought to evaluate if the absence of base editing with shorter gRNAs is due to absence of DNA nicking. We therefore tested the InDel efficiency of gRNAs from 15 to 10 nucleotides length and compared it with full length gRNA in Cas9 stable cells. As expected, there was no InDel formation with any gRNAs except the full-length gRNA (**Supplementary Figure 1A**), suggesting that nicking is absent even at 15 nucleotide length. Additionally, we also tested if any of these gRNAs generated 4.9kb large deletion, which can occur following nuclease activity mediated cleavage at the highly homologous *HBG1* and *HBG2* site. Only full-length gRNA generated large deletions suggesting that nuclease activity is abolished by gRNA length of 15 nucleotides or less (**Supplementary Figure 1B**). However, since HBG_G2_15 even in the absence of nicking showed base conversion, we believe that the loss of base editing in HBG_G2_14 and below is probably due to other factors, probably the insufficient DNA unwinding for deaminase access to the target bases. Additionally, we had previously reported that ABE8e can edit efficiently even in the absence of DNA nicking with full length gRNAs suggesting that the loss of editing observed here might not be related to absence of nicking(George *et al*., 2024).

To validate our findings, we examined truncated variants of a published gRNA targeting *CD33*, a clinically relevant locus in acute myeloid leukemia whose disruption can protect healthy hematopoietic stem cells during CD33-directed therapies(Borot *et al*., 2025). As the gRNA had a monoallelic single nucleotide mismatch at the 12^th^ position in HUDEP-2 cells (**Supplementary Figure 1C**), we first tested if the selected tru-gRNAs can edit at the target site using Cas9, followed by evaluation of base editing efficiency with ABE8e. On editing with Cas9, we noticed that with gRNA truncation, the mismatch tolerance increased, resulting in higher InDel percentage with decreasing lengths up to 17 nucleotides (**Supplementary Figure 1D**). The InDel fraction was close to 100% with CD33_17 gRNA. However, gRNAs CD33_16 and CD33_15 did not show any significant InDel generation, probably because of the loss of catalytic activity. On the other hand, using ABE8e, even with truncation to 15 bp, there was ∼ 60% base conversion(**Figure 1D)**. To our surprise, not only there was no drastic reduction in the editing efficiency in this locus when the gRNAs were truncated, but also there was an increase in the overall editing efficiency at A7 position in all truncated gRNAs compared to the full length gRNA, with ∼ 40% increase in editing efficiency with CD33_17 and CD33_18. It is worth noting that with full length gRNA, the editing at both the target bases were around 50% suggesting that the allele with SNP was not targeted. The increase in editing with truncation could probably be the result of increased mismatch tolerance of shorter gRNAs. However additional studies are required to conclude this observation as truncated gRNAs in general have reduced mismatch tolerance (Lee, Kim and Lee, 2021). This increased mismatch tolerance may be advantageous for rescuing activity at imperfectly matched targets, such as monoallelic SNP-containing loci, but it needs to be evaluated if this may also increase the risk of off-target editing.

Taken together, the data from two different loci suggests that gRNA truncation up to 15 bases can preserve base editing activity although the efficiency varies with different lengths and target context. Our observations also suggest that truncated gRNAs may be useful in regions containing monoallelic SNPs, where they may enable efficient targeting of shared surrounding bases on both alleles.

### Truncated gRNAs can modulate bystander edits in a target dependent manner

One of the major motivations to test truncated gRNAs was to see if they can overcome bystander edits which is a major drawback in base editing especially in coding regions. Based on our observation in *HBG* and *CD33* loci that different bases are edited with substantially different efficiencies by gRNA truncation, we sought to evaluate if this can be applied systematically, to overcome bystander editing. Before targeting specific loci, to evaluate the broader scope, we selected a gRNA in the AAVS1 locus which has multiple adenines in the PAM distal region (**Supplementary Figure 2A**). Base conversion efficiency was evaluated after editing with serially truncated gRNAs. Surprisingly, in this region gRNAs of length 15-17 showed only minimal editing at the target bases (**Figure 2A**). In contrast, AAVS1_18 – AAVS1_20, yielded substantial editing at three positions within the editing window, but with distinct position-specific patterns. While the editing at A7 was similar for all three gRNAs, the editing at A4 was substantially lower for AAVS1_18 and at A6 editing was higher for AAVS1_19. In addition to AAVS1 locus, we selected a gRNA with multiple adenines in the CCR5, a clinically relevant HIV-1 resistance locus (Khan, Paneerselvam and Lawson, 2023), and similarly tested the editing efficiency with ABE8e (**Supplementary Figure 2B**). In this region the overall editing was comparable to AAVS1 locus with, CCR5_16 and CCR5_17 showing minimal editing and a drastic increase in editing with gRNA lengths 18 and above (**Figure 2B**). Additionally, in both the loci, 19 nucleotide gRNAs produced the highest overall base conversion. In CCR5 locus, the highest editing frequency was obtained at A6 position with all lengths of gRNAs and the editing at the flanking bases on both sides dropped in a gRNA dependent manner but following the overall trend. An important observation here is at the A3 position, which when became the 5’ end of gRNA in CCR5_18, showed minimal editing. When plotted as the ratio of editing over the maximum edited base, we can note that the editing of bases at the 5’ end decreases with truncation (**Figure 2C-D**).

**Figure 2:**
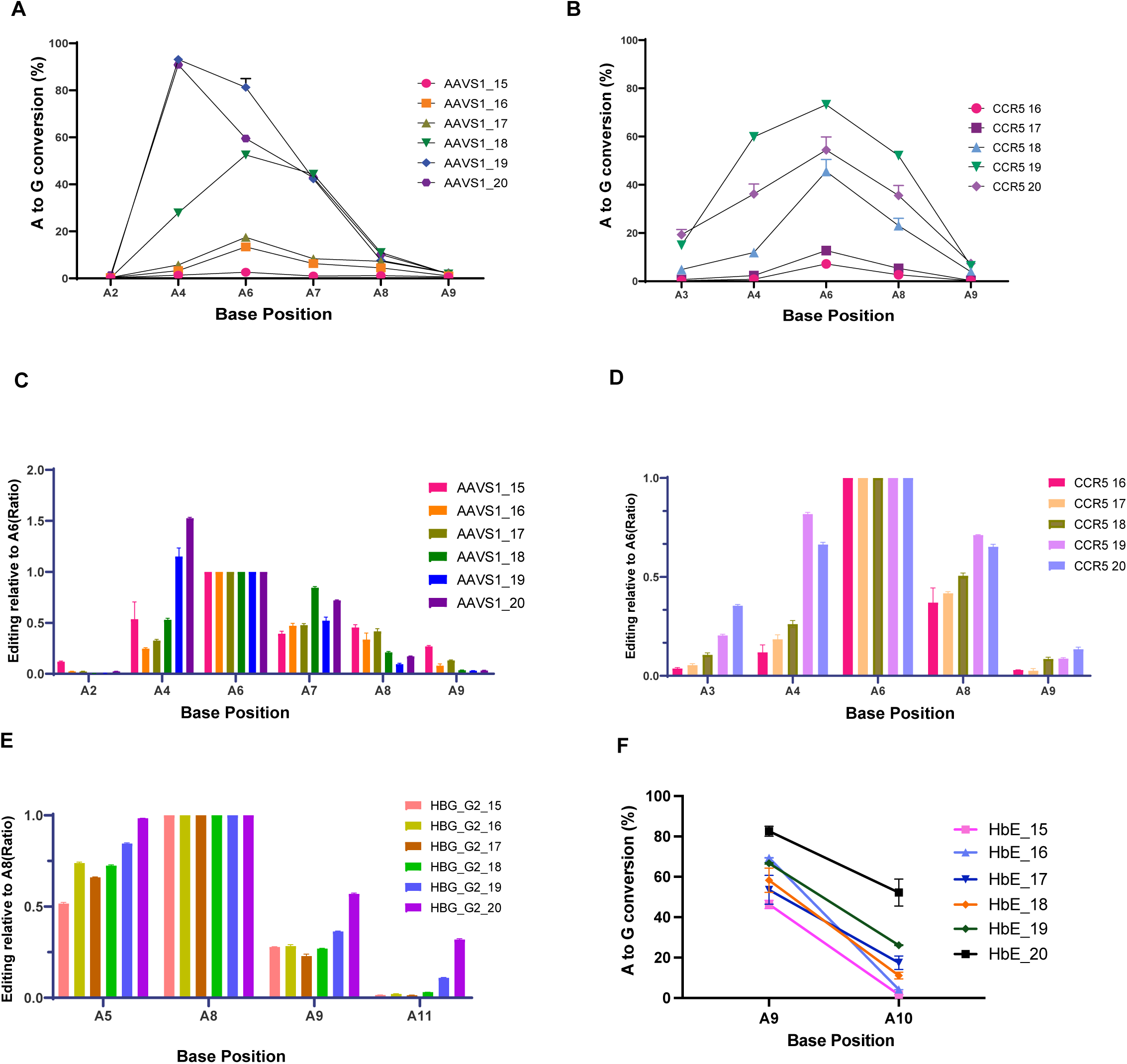
Truncated gRNAs can reduce bystander editing in a locus dependent manner. A) Base conversion efficiency at different positions for 5’truncated gRNAs using ABE8e in the AAVS1 locus by lentiviral delivery B) Base conversion efficiency at different positions for 5’truncated gRNAs using ABE8e in the CCR5 locus by lentiviral delivery C) Editing at various base positions in the AAVS1 locus compared to the maximally edited base D) Editing at various base positions in the CCR5 locus compared to the maximally edited base E) Editing at various base positions in the gamma globin locus compared to the maximally edited base F) Base conversion efficiency at A9 and A10 positions for 5’truncated gRNAs using ABE8e in the Beta globin locus for correction of HbE mutation by lentiviral delivery Data from three independent replicates are plotted as mean± SD.

In the gamma globin promoter as seen before (**Figure 1B**), there was serial variation in editing efficiency with varying gRNA lengths. At A8, which was converted most efficiently in all gRNAs, there was a sudden increase in editing efficiency when gRNA length increased to 16 from 15 and all the incremental lengths showed similar base conversion efficiency. Surprisingly, in addition to A5, the editing at A9 and A11 which are in the 3’ region was also affected significantly by truncation of gRNA. Compared to A8 position the drop in editing was significant at A9 and A11 (**Figure 2E**). These bases correspond to –110 and –112 positions in the gamma globin promoter, which is adjacent to the canonical BCL11A binding site which is covered by A5 and A8(Ravi *et al*., 2022). This reduction in base editing ratios of adjacent bases with gRNA truncations has implications for basic biology application when trying to decode the significance of each bases.

Motivated by the above results and noticing the reduced editing in the 3’ region of editing window, we assessed truncated gRNAs of different lengths for their ability to correct the therapeutically relevant HbE mutation in the β-globin locus using ABE8e(Prasad *et al*., 2024). The pathogenic A9 (HbE) is the desired target mutation while the bystander A>G conversion at A10 creates an undesired haemoglobin variant Hb Aubenas (normal phenotype in heterozygous condition), so it is therapeutically relevant to maximise the editing at A9 while keeping A10 editing to the lowest (**Supplementary Figure 2C**). Interestingly, in this target, although HbE_20 showed the highest overall editing, the truncated HbE_15 gRNA showed the highest on target editing with the least bystander editing bringing the bystander to on target ratio close to 0.03 (**Figure 2F and Supplementary Figure 2D**). Since unbiased genome-wide assessment of off-target activity remains technically challenging for catalytically inactive truncated gRNAs that lack detectable nuclease activity, we performed targeted deep sequencing of the five highest-ranked computationally predicted off-target sites for the HbE_20 gRNA. Comparison of editing outcomes between the full-length HbE_20 gRNA and the truncated HbE_15 gRNA revealed variable effects on off-target editing. OT3 exhibited increased editing with HbE_15 compared with the full-length guide, whereas OT2 and OT4 showed higher editing with HbE_20. At the remaining two sites, editing frequencies were indistinguishable from the negative control (**Supplementary Figure 2E**).

Overall, data from multiple loci indicate that careful gRNA-length selection can reduce, and in some cases abolish, bystander editing while preserving on-target conversion. The effects are target dependent and position specific, so empirical testing will be required to apply this approach broadly.

### Truncated gRNAs can work with other base editors as well

To broaden the applicability, we tested if truncated gRNAs can be used efficiently with other base editors as well. First, we targeted the BCL11A binding site in gamma globin promoter using Cytosine base editor (CBE6b) (Zhang *et al*., 2024). Two gRNAs HBG_G2_20 and HBG_G2_15 were tested in HUDEP-2 cells stably expressing CBE. While the full-length gRNA showed ∼75% editing of both cytosines in the editing window, with the tru-gRNA editing at C6 was reduced to 11% while that at C7 was 42%, bringing the ratio of C6/C7 editing down (**Figure 3 A**). Although we did not test tru-gRNAs of other lengths, we believe CBE should be able to edit with intermediate lengths as HBG_G2_15 edited with similar efficiency as seen with ABE8e.

**Figure 3:**
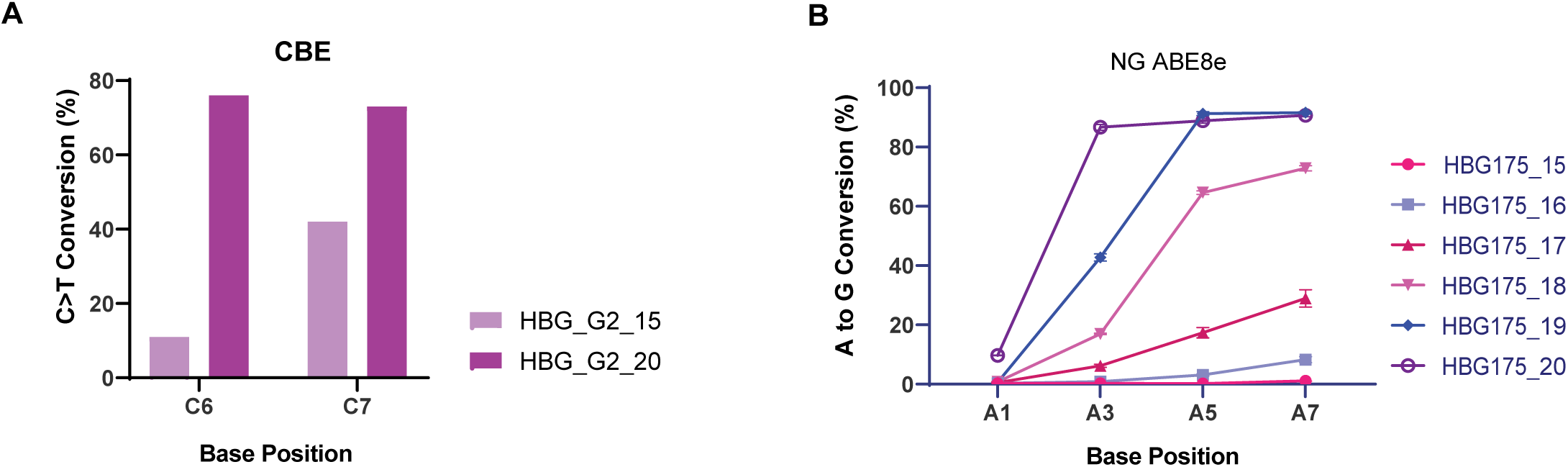
Truncated sgRNAs can edit with different base editors. A) Base conversion efficiency at different positions for full length and 5’truncated gRNA (HBG_G2_15) using TAD CBE in the gamma globin locus by lentiviral delivery B) Base conversion efficiency at different positions for 5’truncated gRNAs using ABE8e NG (NG PAM) in the gamma globin locus targeting the –175 region by lentiviral delivery. Data from three independent replicates are plotted as mean± SD.

Additionally, we tested truncated gRNAs with ABE8e NG (Richter *et al*., 2020) in the gamma globin promoter with the aim to create –175 T>C conversion (HPFH mutation that elevates gamma globin expression by creating a TAL-1 binding site) with minimal conversion at the adjacent bases(Wienert *et al*., 2018). Among the 6 lengths tested, there was no significant editing with HBG175_15 and HBG175_16, but other gRNAs gave measurable editing at all A’s in the editing window. Similar to ABE8e NGG, we noticed a drop in the bystander editing (A3) with HBG175_19 and HBG175_18 compared to full length gRNA (**Figure 3B**). However, with this gRNA we could not variably edit A5 (bystander) and A7 (on target) positions. As was observed at other loci with ABE8e NG, the absence of editing with HBG175_15 and HBG175_16 maybe gRNA specific and validation with additional gRNAs/target sites would be required to conclude if shorter truncated gRNAs are generally compatible with NG-ABE8e.

Overall, the data from different base editors, gRNAs of different lengths and different target loci shows that meticulous selection of gRNA length and the base editor variant can modulate the editing efficiency at a single nucleotide level even to the extent of achieving editing at target base with virtually no editing at the adjacent base.

### Truncated gRNAs can reduce base editing associated large deletions in the globin locus

Base editing is shown to cause large deletions in the globin locus due to simultaneous binding of gRNAs and concurrent nicking at the homologous region(Ravi *et al*., 2022; George *et al*., 2024; Prasad *et al*., 2024). We tested if the frequency of these large deletions is reduced with reducing gRNA length as tru-gRNAs of less than 15 nucleotide is known to lose the cleavage activity. We observed that the large deletions are overcome by the tru-gRNAs of 15 nucleotide length, at both gamma globin (–115 site) and beta globin target sites (HbE mutation) (Supplementary **Figure 3A and B**). With ABE8e, although the deletion frequency was minimal even with full length gRNA, there was a total absence of deletion when the gRNA length was truncated to 16 nucleotides corresponding to the loss of nicking activity. This is similar to the observations with the use of deadCas9 ABE8e where we showed that abolishing the nickase activity can overcome large deletions (George *et al*., 2024). However, it is noteworthy that the editing efficiency was much higher with the use of dABE8e than while using tru-gRNA of 16 nucleotide length. Hence, if the objective is to abolish large deletions in base editing it would be worthwhile to use dABE8e approach rather than truncated gRNAs.

### Truncated gRNAs edit with transient delivery and in primary cells

The previous experiments were performed in stable cell lines with lentiviral delivery of gRNAs, but often disease modelling and therapeutic gene editing requires delivery of components transiently. Hence, we sought to evaluate if the efficiency of truncated gRNAs would vary when delivered transiently as RNA. mRNA encoding the base editors along with 15–20 nucleotide gRNAs were electroporated into HUDEP2 cells. In contrast to lenti viral delivery, the gRNAs targeting gamma globin locus (HBG_G2) did not show editing in either ABE8e or CBE TAD with HBG_G2_15 and HBG_G2_16. The gRNAs above 17 nucleotide length showed editing comparable to lenti viral editing(**Figure 4A and B**). With HbE gRNA the editing efficiency with 15 and 16 nucleotides was very low but measurable and other gRNAs (17-20) showed comparable editing at A9. In contrast to lentiviral editing, the ratio of A10/A9 editing varied significantly with transient delivery in this region with HbE_17 showing comparable editing to HbE_20 at A9 position and virtually absent base conversion at A10 position (**Figure 4C**). The variations in bystander editing was evident in both ABE8e and CBE and at both loci highlighting the potential of using truncated gRNAs for disease modelling or elucidating the role of specific nucleotides in cell lines.

**Figure 4:**
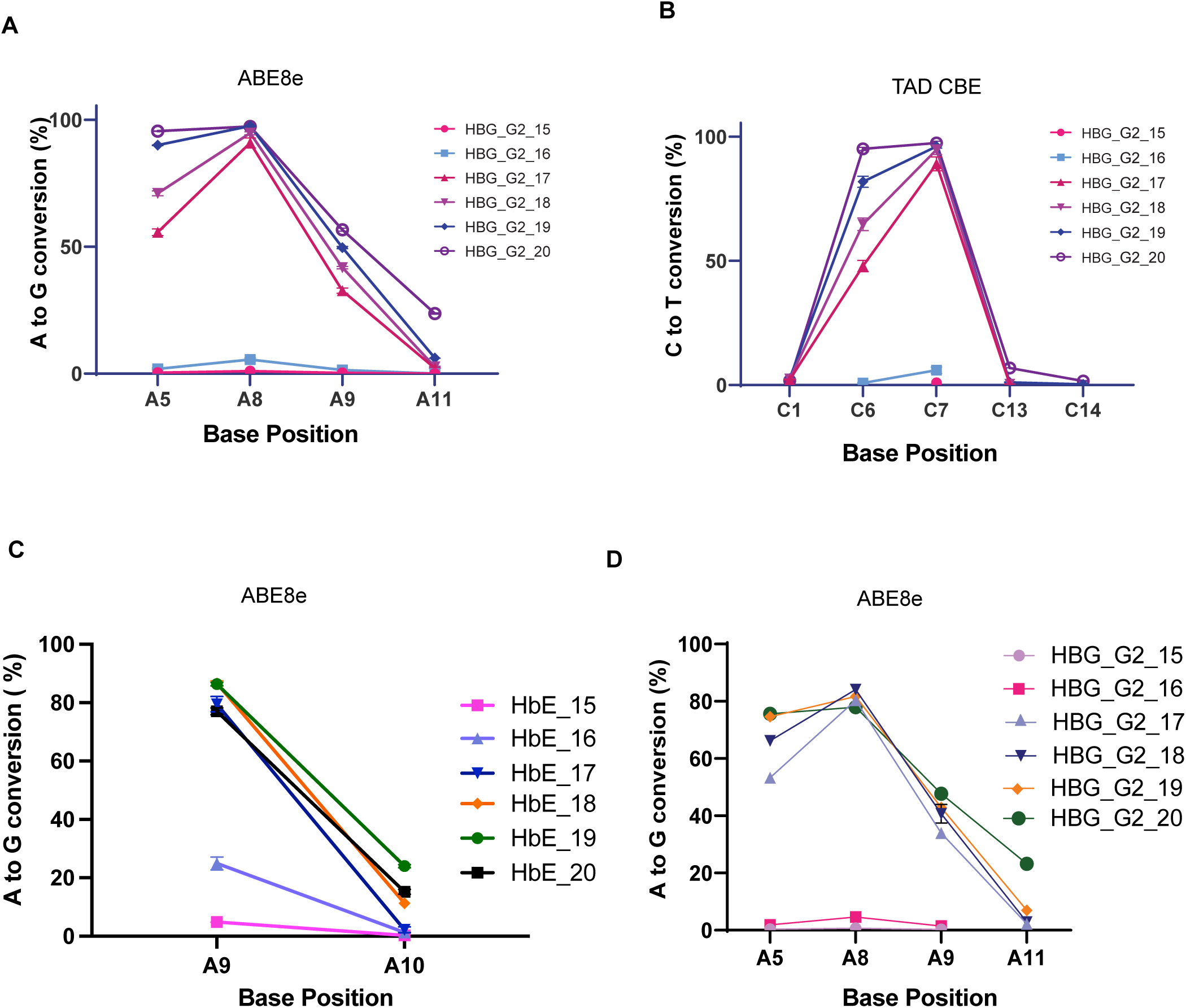
Truncated gRNAs are capable of base editing even with transient delivery. A) Base conversion efficiency at different positions for 5’truncated gRNAs using ABE8e in the gamma globin locus in HUDEP-2 cells upon transient delivery of editing components by nucleofection B) Base conversion efficiency at different positions for 5’truncated gRNAs using TAD CBE in the gamma globin locus in HUDEP-2 cells upon transient delivery of editing components by nucleofection C) Base conversion efficiency at different positions for 5’truncated gRNAs using ABE8e in the beta globin locus in HUDEP-2 cells upon transient delivery of editing components by nucleofection D) Base conversion efficiency at different positions for 5’truncated gRNAs using ABE8e in the gamma globin locus in HSPCs upon transient delivery of editing components by nucleofection Data from three independent replicates are plotted as mean± SD.

In addition to cell lines, for therapeutic editing it is important to be able to edit in primary cells hence we tested the efficiency of HBG_G2 truncated gRNAs in HSPCs with ABE8e. ABE8e was delivered as mRNA and truncated gRNAs were delivered as synthetic RNAs. We observed results comparable to transient cell line editing, in HSPCs as well with A8 position giving the maximum editing. HBG_G2_15 and HBG_G2_16 did not edit in HSPCs either (**Figure 4D**). These findings suggest that transient delivery of base editors favours longer tru-gRNAs (≥17 nucleotide) for efficient on-target editing but testing in multiple loci is required to reach a conclusion.

## Discussion

Our study systematically investigated the impact of gRNA truncation on base editing efficiency and the modulation of bystander editing across multiple genomic loci. Although tru-gRNAs have previously been used in gene editing, to our knowledge no prior work has systematically assessed how truncation affects editing at single-base resolution. Our findings, combined with existing literature, reveal several key insights.

Similar to the reports that Cas9 loses nuclease activity with ≤15 nucleotide gRNAs, we observed absence of base editing at gRNA lengths <15. However, we believe this loss of base editing is due to the loss of accessibility of deaminase to the target site rather than due to the absence of nicking. This is evidenced by the presence of significant editing even with very low nuclease activity with truncated gRNAs targeting *HBG* promoter as well as CD33 locus. We had also demonstrated previously that nicking is dispensable for the activity of newer efficient versions of base editors.

One of the major observation is that tru-gRNAs show strong target-dependent variability in editing efficiency and modulation of bystander edits. We believe that more than truncation, this could be a reflection of factors contributing to DNA unwinding including the sequence composition of gRNA and the local chromatin features(Zhang *et al*., 2016; Cofsky *et al*., 2022; Sinan and Russell, 2022; Li *et al*., 2023). One surprising observation is the variation in editing efficiency at both 5’ and 3’ bases flanking the maximally edited site. Although large scale experiments might be able to shed light on general features guiding the tru-gRNA efficiency, our observations suggest that it is worthwhile to test gRNAs of varying lengths if on-target or bystander editing modulation is intended.

Beyond improving editing precision, tru-gRNAs have broad potential applications in both basic research and therapeutic genome editing. In basic science, the ability to selectively modulate editing at individual nucleotides within the editing window provides a valuable tool for dissecting the functional contribution of closely spaced bases in regulatory elements, promoters, enhancers, and transcription factor binding sites without the need to engineer new base editor variants. Such an approach could facilitate systematic interrogation of cis-regulatory sequences and enable more precise disease modelling by generating defined nucleotide combinations while minimizing confounding bystander edits.

From a therapeutic perspective, gRNA truncation offers a simple and readily implementable strategy to optimize editing outcomes for individual targets, complementing advances in base editor engineering rather than replacing them. Because gRNA length can be modified without altering the editing enzyme or delivery platform, it provides an additional layer of optimization that can be incorporated into existing therapeutic workflows.

A limitation of this study is that we were unable to perform a comprehensive assessment of off-target editing induced by tru-gRNAs. Although comprehensive genome-wide off-target analysis would strengthen the study, our primary objective was to determine how guide truncation influences on-target editing efficiency and editing-window composition. Existing genome-wide off-target detection methods are largely optimized for nuclease-mediated DNA double-strand breaks and are therefore not readily applicable to tru-gRNAs, which lack detectable nuclease activity(Guo *et al*., 2023; Xu *et al*., 2026). Furthermore, gRNA truncation substantially expands the number of potential genomic sites with partial complementarity, making targeted validation based solely on in silico prediction impractical. Development of robust approaches for genome-wide profiling of off-target deaminase activity in the absence of DNA cleavage will be important for future studies.

In conclusion, our work expands the current framework for optimizing base editing by demonstrating that gRNA length can be exploited as an independent design parameter to fine-tune editing outcomes. Rather than functioning as an all-or-none determinant of editing activity, gRNA length reshapes editing efficiency and the editing window in a locus-dependent manner, allowing selective modulation of individual nucleotides without altering the editor itself. Across multiple loci, base editors, cell types and delivery platforms, tru-gRNAs retained measurable editing activity while altering editing patterns. Although comprehensive characterization across additional genomic contexts and assessment of off-target effects will be required before broad application, these findings provide a practical and easily implementable strategy that complements advances in base editor engineering and may facilitate more precise genome editing for both research and therapeutic applications.

## Data availability statement

All data supporting the findings of this study are available in the main text and Supplementary information. Additional supporting data are available from the corresponding author upon reasonable request.

## Author contributions

AG—Conceptualization, methodology, formal analysis, data curation, visualisation, writing-original draft; KP – Methodology, formal analysis, data curation, visualisation writing-review and editing; PS – Methodology, data curation, writing-review and editing; KMM-Supervision, formal analysis, funding acquisition, Conceptualization, Investigation, visualisation, project administration, resources, Writing-review and editing; SM,ST – resources, Writing-review and editing. All authors read and approved the submitted version.

## Acknowledgments

We thank all the members of Advanced Genome Editing Lab, CSCR for their valuable inputs. We acknowledge the CSCR Core for providing access to instrumentation and for their support for flow cytometry experiments.

