## Supplementary table and figures for "Guide RNA Truncation as a Tunable Approach for Modulating Base Editing Outcomes"

### Supplementary Figure 1

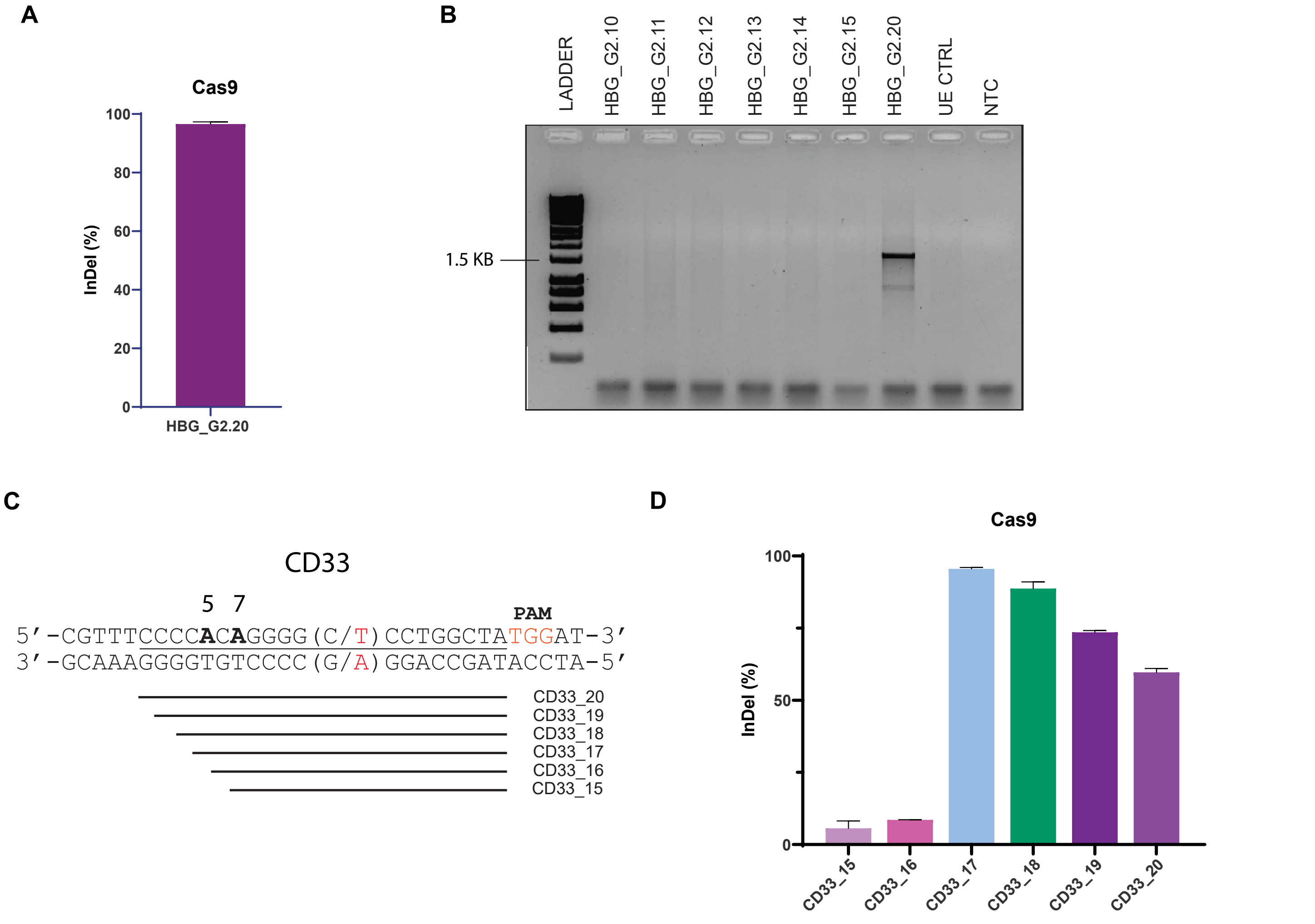


**Supplementary Figure 1 : Cas9 mediated gene editing using truncated gRNAs**

A) InDel frequency at the gamma globin locus on editing with Cas9 and full length gRNA.
B) Formation of fusion gene (HBG2 promoter -HBG1) due to large intergenic deletion detected by agarose gel electrophoresis following PCR. A band at ~1700 bp indicates the amplicons from fusion gene.
C) Sequence of full length and truncated gRNAs targeting the CD33 locus. Target Adenosines within the editing window are represented in bold letters and the monoallelic SNP is represented in the brackets.
D) InDel frequency at the CD33 locus on editing with Cas9 and truncated gRNAs.

#### Supplementary Figure 2

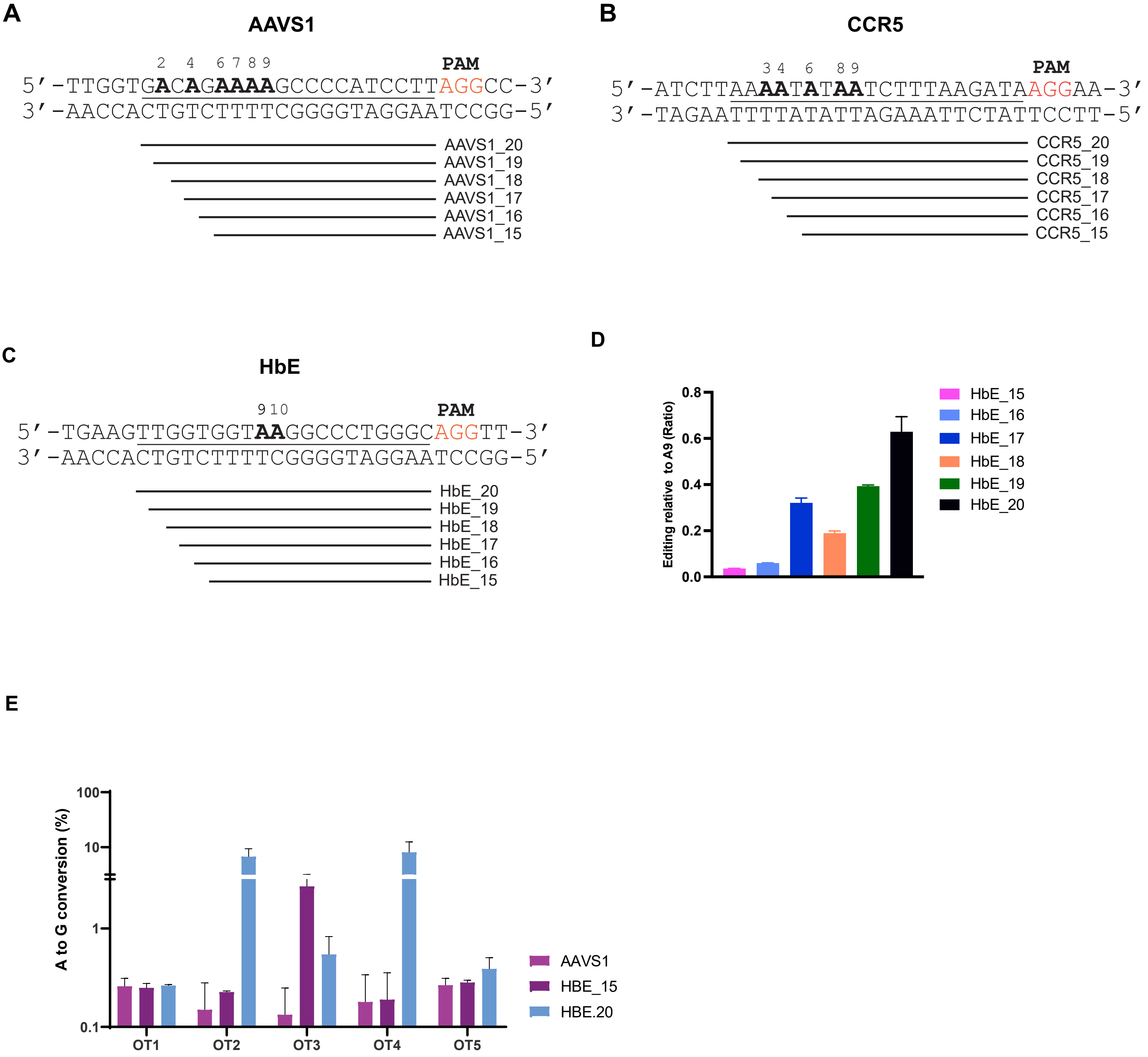


**Supplementary Figure 2: Targeting different loci using truncated sgRNAs**

A) Sequence of full length and truncated gRNAs targeting the AAVS1 locus. Target Adenosines within the editing window are represented in bold letters

B) Sequence of full length and truncated gRNAs targeting the CCR5 locus. Target Adenosines within the editing window are represented in bold letters

C) Sequence of full length and truncated gRNAs targeting the HBB locus (HbE gRNA). Target Adenosines within the editing window are represented in bold letters

D) Editing at A10 position (bystander editing) compared to A9 (on-target) in the HbE gRNA

E) Off-target base conversion at the top 5 sites for HbE gRNA (full length) predicted by in-silico tools. Base conversion at the same sites in cells edited with AAVS1 gRNA was used as control.

**Supplementary Figure 3**

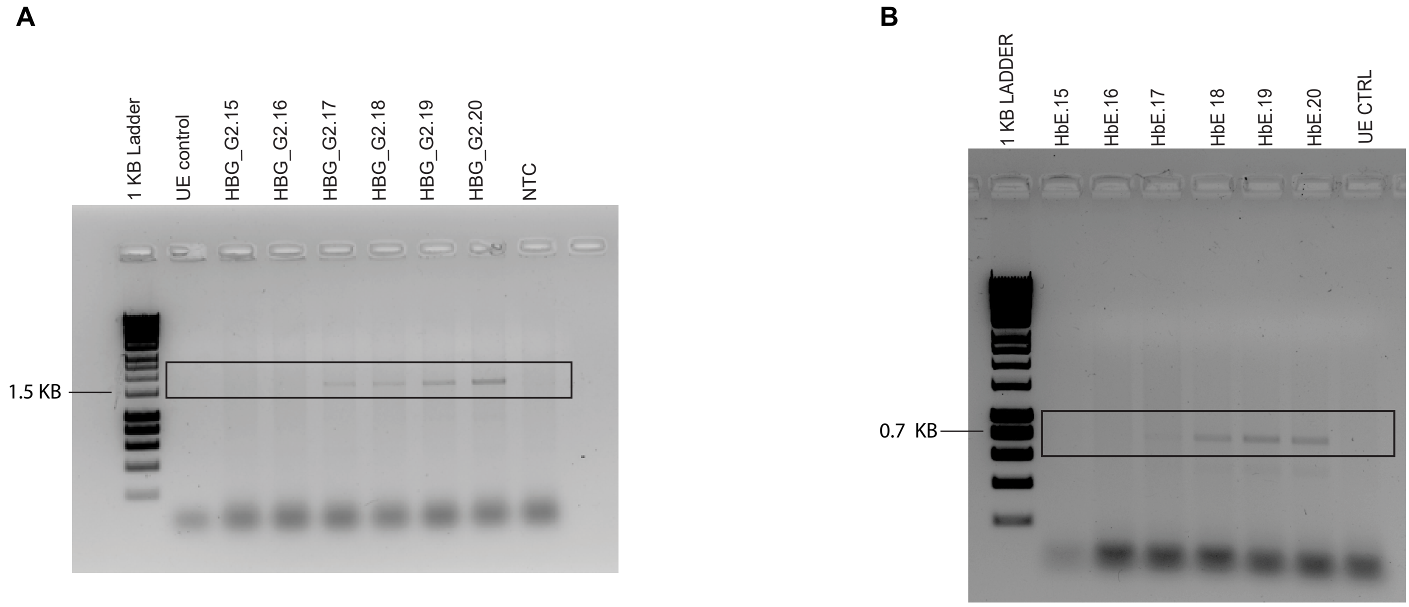


**Supplementary Figure 3: Simultaneous nick mediated large deletions can be reduced by truncation of gRNAs**

A) Formation of fusion gene (HBG2 promoter -HBG1) due to large intergenic deletion detected by agarose gel electrophoresis following PCR. A band at ~1700 bp indicates the amplicons from fusion gene. Data from HUDEP-2 cells edited by ABE8e via nucleofection.

B) Formation of fusion gene (HBD-HBB) due to large intergenic deletion detected by agarose gel electrophoresis following PCR. A band at ~700 bp indicates the amplicons from fusion gene. Data from HbBE HUDEP-2 cells edited by ABE8e via nucleofection.

### List of gRNAs and primers used in the study

**Supplementary Table 1: List of gRNAs used in the study**

| Sl. No. | Name | Sequence (5'- 3’) |
|---|---|---|
| 1. | HBG_G2_10 | AGCCTTGACA |
| 2. | HBG_G2_11 | TAGCCTTGACA |
| 3. | HBG_G2_12 | ATAGCCTTGACA |
| 4. | HBG_G2_13 | AATAGCCTTGACA |
| 5. | HBG_G2_14 | CAATAGCCTTGACA |
| 6. | HBG_G2_15 | CCAATAGCCTTGACA |
| 7. | HBG_G2_16 | ACCAATAGCCTTGACA |
| 8. | HBG_G2_17 | GACCAATAGCCTTGACA |
| 9. | HBG_G2_18 | TGACCAATAGCCTTGACA |
| 10. | HBG_G2_19 | TTGACCAATAGCCTTGACA |
| 11. | HBG_G2_20 | CTTGACCAATAGCCTTGACA |
| 12. | HBG_175_15_NG | TATTTGCATTGAGAT |
| 13. | HBG_175_16_NG | ATATTTGCATTGAGAT |
| 14. | HBG_175_17_NG | GATATTTGCATTGAGAT |
| 15. | HBG_175_18_NG | AGATATTTGCATTGAGAT |
| 16. | HBG_175_19_NG | CAGATATTTGCATTGAGAT |
| 17. | HBG_175_20_NG | ACAGATATTTGCATTGAGAT |
| 18. | CCR5_16 | TATAATCTTTAAGATA |
| 19. | CCR5_17 | ATATAATCTTTAAGATA |
| 20. | CCR5_18 | AATATAATCTTTAAGATA |
| 21. | CCR5_19 | AAATATAATCTTTAAGATA |
| 22. | CCR5_20 | AAAATATAATCTTTAAGATA |
| 23. | AAVS1_15 | AAAAGCCCCATCCTT |
| 24. | AAVS1_16 | GAAAAGCCCCATCCTT |
| 25. | AAVS1_17 | AGAAAAGCCCCATCCTT |
| 26. | AAVS1_18 | CAGAAAAGCCCCATCCTT |
| 27. | AAVS1_19 | ACAGAAAAGCCCCATCCTT |
| 28. | AAVS1_20 | GACAGAAAAGCCCCATCCTT |
| 29. | CD33_15 | CAGGGGCCCTGGCTA |
| 30. | CD33_16 | ACAGGGGCCCTGGCTA |
| 31. | CD33_17 | CACAGGGGCCCTGGCTA |
| 32. | CD33_18 | CCACAGGGGCCCTGGCTA |
| 33. | CD33_19 | CCCACAGGGGCCCTGGCTA |
| 34. | CD33_20 | CCCCACAGGGGCCCTGGCTA |
| 35. | HbE_15 | GGTAAGGCCCTGGGC |
| 36. | HbE_16 | TGGTAAGGCCCTGGGC |
| 37. | HbE_17 | GTGGTAAGGCCCTGGGC |
| 38. | HbE_18 | GGTGGTAAGGCCCTGGGC |
| 39. | HbE_19 | TGGTGGTAAGGCCCTGGGC |
| 40. | HbE_20 | TTGGTGGTAAGGCCCTGGGC |

### List of gRNAs and primers used in the study

**Supplementary Table 2: List of primers used in the study**

| Sl. No. | Name | Forward Primer (5' - 3') | Reverse Primer (5' - 3') |
|---|---|---|---|
| 1 | HBG_NGS | TACACGACGCTCTTCCGATCTGGAATGA CTGAATCG GAACAA | AGACGTGTGCTCTTCCGATCTGACGTTCCA GAAGCG AGTGT |
| 2 | AAVS1_NGS | TACACGACGCTCTTCCGATCTAGGTTCT GGCAAAGGAGAGAGATG | AGACGTGTGCTCTTCCGATCTCTATGTCCA CTTCAGGACAGCATG |
| 3 | CCR5_NGS | TACACGACGCTCTTCCGATCTGCTTGGC CTGTTAGTTAGCT | AGACGTGTGCTCTTCCGATCTGCTCCCTTT AAAGTCTTTCA |
| 4 | CD33_NGS | TACACGACGCTCTTCCGATCTTGCTTCC TCAGACATGCCGC | AGACGTGTGCTCTTCCGATCTTGGGATGGA AGAAAGTGCAG |
| 5 | HbE-NGS | TACACGACGCTCTTCCGATCTGCTTCT GACACAACTGTGTT | AGACGTGTGCTCTTCCGATCTTCTTCT CTGTCTCCACATGC |
| 6 | HBG_Del | CCATAGTATCTGGTAAAGAGCA | GATAGATGTTTTAAAACAAC |
| 7 | HBB-HBD_Del | ATGTAGAGGAGAACAGGGTT | CTGTACCCTGTTACTTATCC |
| 8 | HbE- OT1_NGS (Intergenic) Chr19:30679326-30679348 | TACACGACGCTCTTCCGATCTTTAAAGA GCGTTAATGAGCTTCTCAGAACCC | AGACGTGTGCTCTTCCGATCTAGGGGGGT GGCCGTGATGGCAAGA |
| 9 | HbE- OT2_NGS ChrX:15060055-15060077 | TACACGACGCTCTTCCGATCTCAGCAG CCACCGACTTCC | AGACGTGTGCTCTTCCGATCTAAATAATTAC AGTCAGTTTCACTTAAGGGGGAGATC |
| 10 | HbE- OT3_NGS (RP11-30M7) Chr2:234225153-234225175 | TACACGACGCTCTTCCGATCTGGAAGG GTCACAATGTCCACCTTT | AGACGTGTGCTCTTCCGATCTCACTTTACC ACCTTGAGGAGTAGTTGTTG |
| 11 | HbE- OT4_NGS (TBL1X gene) ChrX:9707424-9707446 | TACACGACGCTCTTCCGATCTAGTTTCT ATTAATCATCATGGCACCTCTTGATGG | AGACGTGTGCTCTTCCGATCTCCAACAACT TTGTCTTTTTTACACCGTAAAATAGCC |
| 12 | HbE- OT5_NGS Chr17:42172056-42172078 | TACACGACGCTCTTCCGATCTTTTCTCC TTACAACCTCCCAATCTACCC | AGACGTGTGCTCTTCCGATCTGTGCAGGG AAGTCCTCTTGG |
